# Assessment of an AI virtual staining model performance across same and serial tissue sections using CD3^+^ T cell ground truth

**DOI:** 10.1101/2023.11.12.565422

**Authors:** Abu Bakr Azam, Felicia Wee, Juha P. Väyrynen, Willa Wen-You Yim, Yue Zhen Xue, Bok Leong Chua, Jeffrey Chun Tatt Lim, Daniel Shao Weng Tan, Angela Takano, Chun Yuen Chow, Li Yan Khor, Tony Kiat Hon Lim, Joe Yeong, Mai Chan Lau, Yiyu Cai

## Abstract

Immunophenotyping via multi-marker assays significantly contributes to patient selection, therapeutic monitoring, biomarker discovery, and personalized treatments. Despite its potential, the multiplex immunofluorescence (mIF) technique faces adoption challenges due to technical and financial constraints. Alternatively, hematoxylin and eosin (H&E)-based prediction models of cell phenotypes can provide crucial insights into tumor-immune cell interactions and advance immunotherapy. Current methods mostly rely on manually annotated cell label ground truths, with limitations including high variability and substantial labor costs. To mitigate these issues, researchers are increasingly turning to digitized cell-level data for accurate in-situ cell type prediction. Typically, immunohistochemical (IHC) staining is applied to a tissue section serial to one stained with H&E. However, this method may introduce distortions and tissue section shifts, challenging the assumption of consistent cellular locations. Conversely, mIF overcomes these limitations by allowing for mIF and H&E staining on the same tissue section. Importantly, the multiplexing capability of mIF allows for a thorough analysis of the tumor microenvironment by quantifying multiple cell markers within the same tissue section. In this study, we introduce a Pix2Pix generative adversarial network (P2P-GAN)-based virtual staining model, using CD3^+^ T-cells in lung cancer as a proof-of-concept. Using an independent CD3 IHC-stained lung cohort, we demonstrate that the model trained with cell label ground-truth from the same tissue section as H&E staining performed significantly better in both CD3^+^ and CD3^-^ T-cell prediction. Moreover, the model also displayed prognostic significance on a public lung cohort, demonstrating its potential clinical utility. Notably, our proposed P2P-GAN virtual staining model facilitates image-to-image translation, enabling further spatial analysis of the predicted immune cells, deepening our understanding of tumor-immune interactions, and propelling advancements in personalized immunotherapy. This concept holds potential for the prediction of other cell phenotypes, including CD4^+^, CD8^+^, and CD20^+^ cells.

## Introduction

Immune phenotyping in tissue, facilitated by multi-marker assays such as mIF, plays a pivotal role in patient selection, treatment monitoring, biomarker discovery, and the development of targeted and personalized therapeutic strategies^1,2,3^. Nevertheless, the wider adoption of the mIF technique faces challenges as it remains inaccessible to many laboratories due to technical and time constraints or funding limitations. Conversely, the utilization of hematoxylin and eosin (H&E)-based prediction models present a viable alternative for generating data to enhance our comprehension of the intricate interactions within the immune system. Given that H&E staining is cost-effective and routinely performed in numerous histology laboratories, integrating H&E-based prediction models into existing workflows can be achieved with relative ease. This approach has the potential to revolutionize the field of immunotherapy, opening new avenues for advancements in treatment strategies.

Current studies of H&E-based approaches largely rely on manual annotated cell label ground truth^4,5^. For instance, a study by Wilde *et al.* demonstrated the use of deep learning (DL) to assess two prognostic risk parameters, OP-TIL and the multinucleation index (MuNI), in hematoxylin and eosin (H&E) stained slides from patients with oropharyngeal squamous cell carcinoma^6^. The group proposed two DL-based imaging biomarkers, namely OP-TIL, which quantitatively characterizes the spatial patterns between tumor infiltrating lymphocytes (TILs) and their surrounding cells^7^, while the MuNI quantifies the multinucleated tumor cells in epithelial regions^8^. Conditional generative adversarial network (cGAN) models were adopted for cell segmentation based on OP-TIL, and trained for in-silico computation of MuNI. This group also highlighted the potential clinical importance of identification and tissue localization of TIL subtypes, such as those expressing CD4, CD8, and CD20^7^. However, the applicability of these approaches was limited by the availability of manual annotation of TILs and multinucleated tumor cells by pathologists, with high inter- and intra-observer variability and high labor costs^9^.

To address both inter- and intra-observer discrepancies in the annotation and scoring of cell phenotypes, there has been a growing interest in the utilization of digitized cell-level data as the definitive reference for predicting cell types in situ^10^. Commonly, immunohistochemical (IHC) staining is applied to a tissue section that is consecutive to another one stained with H&E, assuming that similar cells maintain identical locations across both sections. Yet, in conventional IHC methods, manual preparation can cause distortions, and heat fixation can shift the tissue section^11^, disrupting this assumption. Furthermore, achieving same-section ground truth is impeded by chromogenic IHC due to deposition of the brown chromogen 3,3’-diaminobenzidine (DAB).

Alternatively, multiplex immunofluorescence (mIF) overcomes these limitations, enabling staining on the same tissue section used for H&E staining. Crucially, mIF’s multiplexing feature allows a comprehensive analysis of the tumor microenvironment (TME) by quantifying multiple cell markers within the same tissue section^12^. In the realm of immunotherapy, the simultaneous quantification of immune markers like CD3, CD4, CD8, cytokeratin, PD-1, and CTLA-4 within the same tissue space is critical for a comprehensive understanding of tumor-immune interactions^13,14^. Here, we propose a Pix2Pix generative adversarial network (P2P-GAN)-based virtual staining model, using CD3^+^ T-cells in lung cancer as the study model (Figure 1a). The choice of CD3^+^ T-cells highlights their significant role in lung cancer prognosis and treatment^15,16,17^. We hypothesize that the performance of the prediction model can be impacted by cellular differences in adjacent (non-identical) tissue sections. To test this hypothesis, we built and compared two DL models, one trained using the CD3^+^ T-cell ground truth obtained by mIF staining of the same tissue section stained with H&E (abbreviated as same-section model; Figure 1b), while the other model was trained using the CD3^+^ T-cell ground truth obtained by mIF staining of the serial tissue section stained with H&E (abbreviated as serial-section model; Figure 1b).

**Figure 1:**
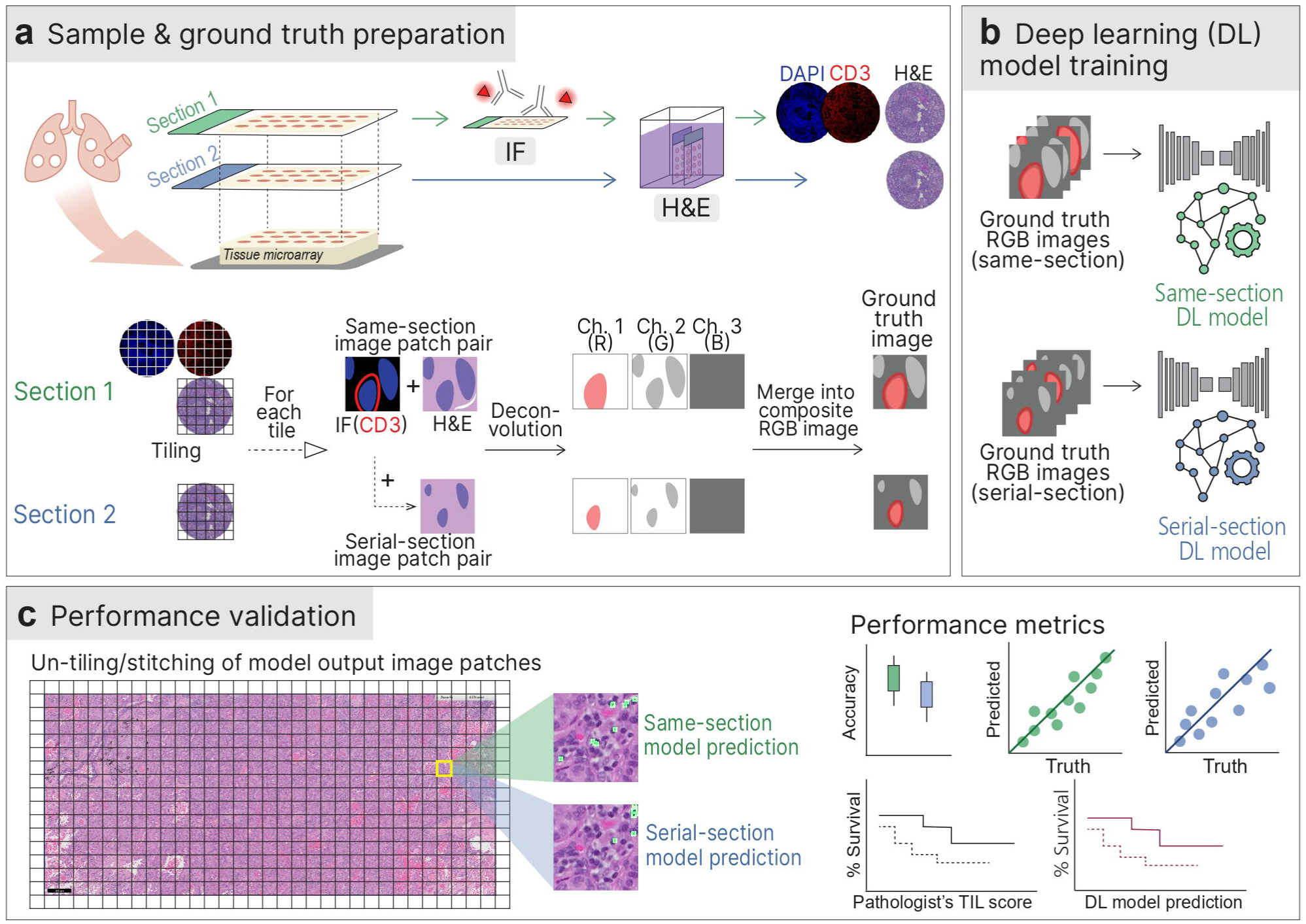
Schematic diagram of the study protocol. a) Preparation of samples and ground truth for both serial-section and same-section datasets; b) Construction and training of two P2P-GAN DL models utilizing the serial-section and same-section datasets; c) Validation of DL model performance using an independent in-house IHC cohort and an external lung cancer cohort.

## Materials/Subjects and Methods

### Cohorts

This study was conducted using three lung cancer cohorts (2 in-house and 1 public). The training cohort consisted of formalin-fixed paraffin embedded (FFPE) tissues in the tissue microarray (TMA) format prepared in the Department of Anatomical Pathology of Singapore General Hospital (Agency of Science, Technology and Research (A*STAR) IRB: 2021-161, 2021-188, 2021-112). The tissue sections were stained with H&E and mIF (anti-CD3 and DAPI for nuclear staining) in the Institute of Molecular and Cell Biology (IMCB) at the Agency for Science, Technology and Research, Singapore. Using this cohort, we prepared the same-section and serial-section datasets. In the same-section dataset, 57 H&E and mIF image pairs were generated from the same tissue sections of the 57 patients. In the serial-section dataset, a separate set of H&E images were generated using tissue sections adjacent to the tissue sections used for the mIF staining.

Separate in-house and public cohorts were used for evaluation of the model performance (Table 1). The in-house cohort comprised CD3 IHC-stained images along with H&E images generated from the corresponding serial-section in TMAs (designated the IHC cohort). The public cohort consisted of H&E-stained images (20× magnification) and the companion patient survival data were downloaded from OncoSG (Singapore Oncology, Data Portal) (designated the Onco-SG cohort).

**Table 1:**
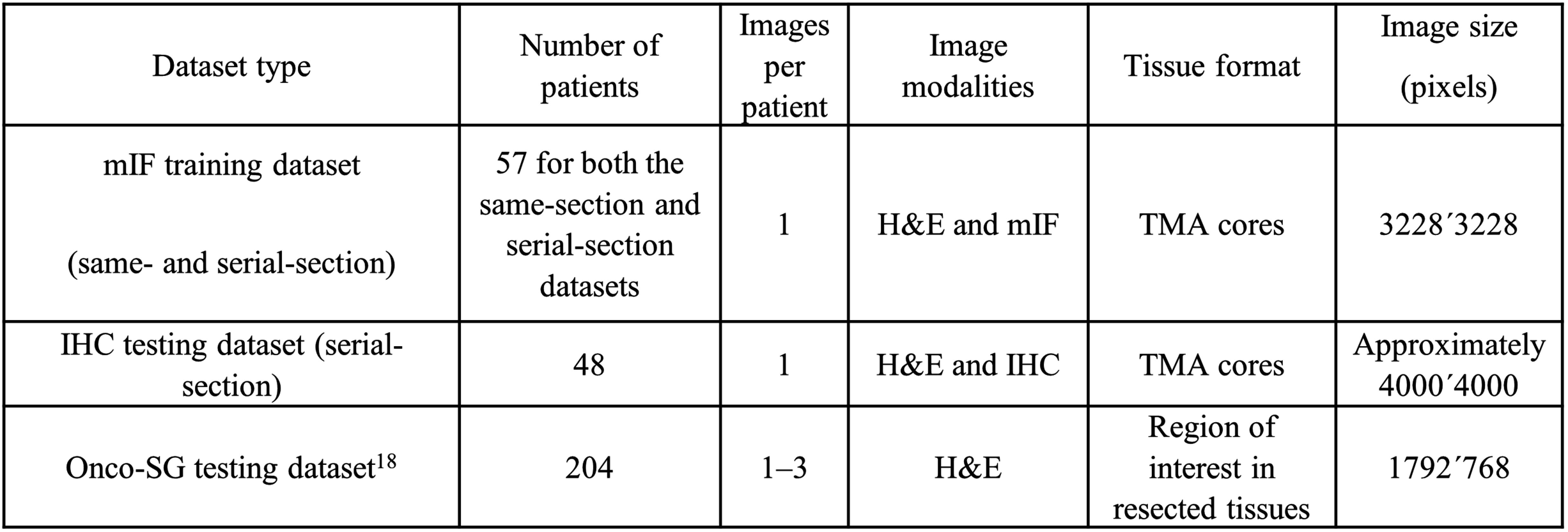
Cohort characteristics.

### Tissue staining

The FFPE tissues were sectioned (4 µm thickness) and heat-fixed at 65°C for 5 min before manual staining with hematoxylin (Epredia, Fisher Scientific, Porto Salvo, Portugal) and eosin (Epredia, Fisher Scientific, Gothenburg, Sweden). IHC staining was performed on the FFPE tissues (4 µm thickness) with anti-CD3 primary antibody (1:200; Dako A0452, Santa Clara, CA, USA) using the Leica Bond Max autostainer (Leica Biosystems, Melbourne, Australia) and Bond Refine Detection Kit (Leica Biosystems) as previously described^19^. The H&E and IHC stained slides were then scanned using the Axioscan.Z1 Slide Scanner (Zeiss, Oberkochen, Germany).

Next, mIF staining was performed on the FFPE tissue sections (4 µm thickness) using the Leica Bond Max autostainer (Leica Biosystems, Melbourne, Australia), Bond Refine Detection Kit (Leica Biosystems) and Opal 6-Plex Detection Kit for Whole Slide Imaging (Akoya Biosciences, Marlborough, MA, USA) as previously described^19^. In brief, FFPE tissue sections were subjected to repeated cycles of heat-induced epitope retrieval, incubation with anti-CD3 primary antibody (Dako #A0452), anti-rabbit poly-HRP-IgG (Ready-to-use; Leica Biosystems) and Opal tyramide signal amplification (TSA) (Akoya Biosciences). Spectral DAPI (4’,6-diamidino-2-phenylindole) (Akoya Biosciences) was applied as the final nuclear counterstain. Images were captured using the Vectra 3 Automated Quantitative Pathology Imaging System (Akoya Biosciences). After scanning, the mIF slides were subjected to H&E staining, followed by scanning on the Axioscan.Z1 Slide Scanner (Zeiss).

### Ground truth cell labels

For model training, ground truth cell labelling involved the identification of CD3^+^ cells in the H&E image space according to a series of steps. First, nuclei in the H&E image were identified using the StarDist Python library (pre-trained for H&E images)^20^. Second, nuclei and CD3^+^ regions in the mIF image were identified individually using the StarDist Python library (pre-trained for fluorescence images) based on DAPI staining and CD3 expression, respectively. These regions were then overlaid to identify CD3^+^ T-cells in the mIF image. Third, the CD3^+^ T-cells identified in the mIF image were matched to the closest nuclei in the H&E image stained on the same (post-mIF H&E staining) or serial tissue section (designated same-section and serial-section datasets, respectively). The H&E image with CD3^+^ information (i.e., ground truth image) was then deconvoluted into red (R), green (G), and blue (B) channels representing the CD3^+^ T-cell, haematoxylin (H), and eosin (E) staining, respectively. Representation of the CD3^+^ T-cell information in a separate channel i.e., R, facilitates the identification of predicted CD3^+^ T-cells during model deployment. Considering that CD3 localizes to the cell membrane whereas DAPI staining is localized in the nucleus, Gaussian noise (kernel size 101) was applied to the R channel of the image to increase the spread of the CD3^+^ signals while keeping the maximum intensity at its center. This facilitates the identification of predicted CD3^+^ T-cells, which relies on an overlap between CD3 and DAPI intensities i.e., R and G channels.

In the IHC testing dataset, CD3 signal localization in an IHC image was first determined by applying a threshold (value >100) to the DAB stain intensity, resulting in a binary mask where 1 indicates CD3 detection and 0 indicates otherwise. The CD3 mask was then overlaid on the nuclei segmented in the paired H&E image to identify CD3^+^ T-cells (ground truth cell labels) according to the same procedure described for mIF dataset. In the Onco-SG testing dataset, two pathologists (YZX and JPV) assessed the H&E images and scored the %TIL. Model performance was evaluated by comparing overall %CD3^+^ T-cell with the %TIL in individual patients by Spearman’s correlation analysis. We also assessed the 5-year overall survival association with the patient groups stratified using the mean DL-predicted %CD3^+^ T-cell versus the mean of the %TIL values determined by the two pathologists. If multiple images were available for the same patient, the patient-average %CD3^+^ T-cell or %TIL value was used.

### P2P-GAN model architecture

A conventional GAN incorporates a generative network to produce image candidates and a discriminative network for their evaluation. The former network is trained to ‘fool’ the latter, hence facilitating unsupervised learning by the model. The P2P-GAN is a variation of a conditional GAN, in which the generator output image is conditional on the input image, and hence is designed perfectly for the image-to-image translation task. In this study, we adopted the P2P-GAN architecture reported by Isola *et al*.^21^ in which a U-Net was used as the generator and a convolutional neural network (CNN) was used as the discriminator (Figure 2). Model training involved presenting the generator with stain-deconvoluted H&E images, while presenting the discriminator with ground truth images (i.e., stain-deconvoluted H&E images overlaid with mIF-identified CD3^+^ T-cell information). These images were then compared with the generator predicted images to output a 30×30 matrix for updating both the generator and discriminator (Figure 2; more details are provided below).

**Figure 2:**
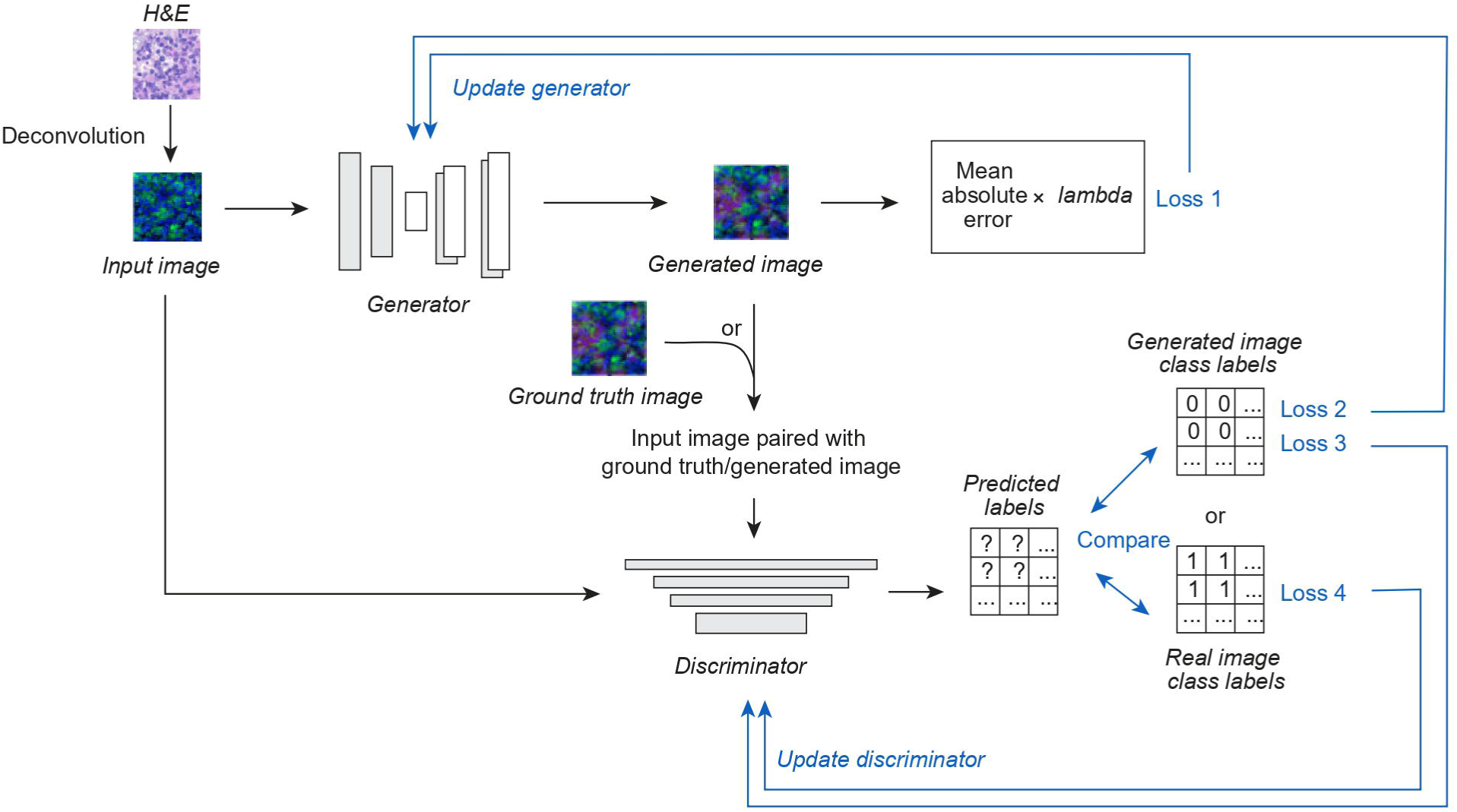
P2P-GAN model architecture and parameter updating process during model training. Two key components, namely the generator, which inputs the H&E image patch and generates (predicts) CD3+ signals on the input image, and its adversary, the discriminator, which distinguishes the generator output from the image with true CD3+ signals (ground truth). The adversarial nature of the network enables the generator to produce good predictions.

### Model training

Two P2P-GAN models were trained using the same-section and serial-section training datasets (henceforth referred to as the same-section and serial-section models, respectively). Each image in the training dataset (Table 1) was divided into 256×256 image patches (total 9,633 patches). Of these, 96% (9,249 patches) were used for model training and 4% (384 patches) were randomly selected for model testing (hereafter referred as the held-out subset). The generator and discriminator work in an adversarial fashion such that the respective losses are balanced out. The overall objective is to reach an optimum for the two conflicting goals, where the generator produces an output that is almost indistinguishable from the ground truth images, while the discriminator can distinguish images generated by the generator from ground truth images. Overall, three different types of losses must be minimized: LOSS 1, which measures the mean absolute difference between the generator output image and the ground truth image, is used to update the generator network; LOSS2/LOSS 3 and LOSS 4 measure the difference between the 30×30 feature matrix output from the discriminator with two 30×30 target matrices, one of which contains all 0 digits and the other contains all 1 digits. This allows quantification of ‘lack of capability’ and ‘capability’, respectively, of the discriminator in distinguishing the generator output image; LOSS2 (essentially LOSS3) is feedback to the generator, while LOSS 3 and LOSS4 are feedback to the discriminator (Figure 2). The training of both models involved 150 epochs with a batch size of 350. A regularization value of 100 was applied to LOSS 1 (i.e., the mean absolute loss).

### Model performance characteristics

Model performance was quantified based on two key metrics, namely CD3^+^ and CD3^-^ T-cell counts, and overall accuracy (defined as the ratio of correctly predicted CD3^+^ and CD3^-^ T-cell counts to the total number of cells). The model-predicted CD3^+^ and CD3^-^ T-cell counts were identified as shown in Figure 3. Specifically, model-predicted CD3 signals (represented in the red channel) were overlaid with the nuclei segmented from the input H&E image to identify the CD3^+^ T-cell, whereas nuclei (or cells) with no matching CD3 signals were deemed to be CD3^-^ T cells. The model-predicted CD3^+^ and CD3^-^ T-cell values were then overlaid with the paired mIF (training cohort) or IHC (testing cohort) images to quantify the accurately predicted CD3^+^ and CD3^-^ T-cell counts.

**Figure 3:**
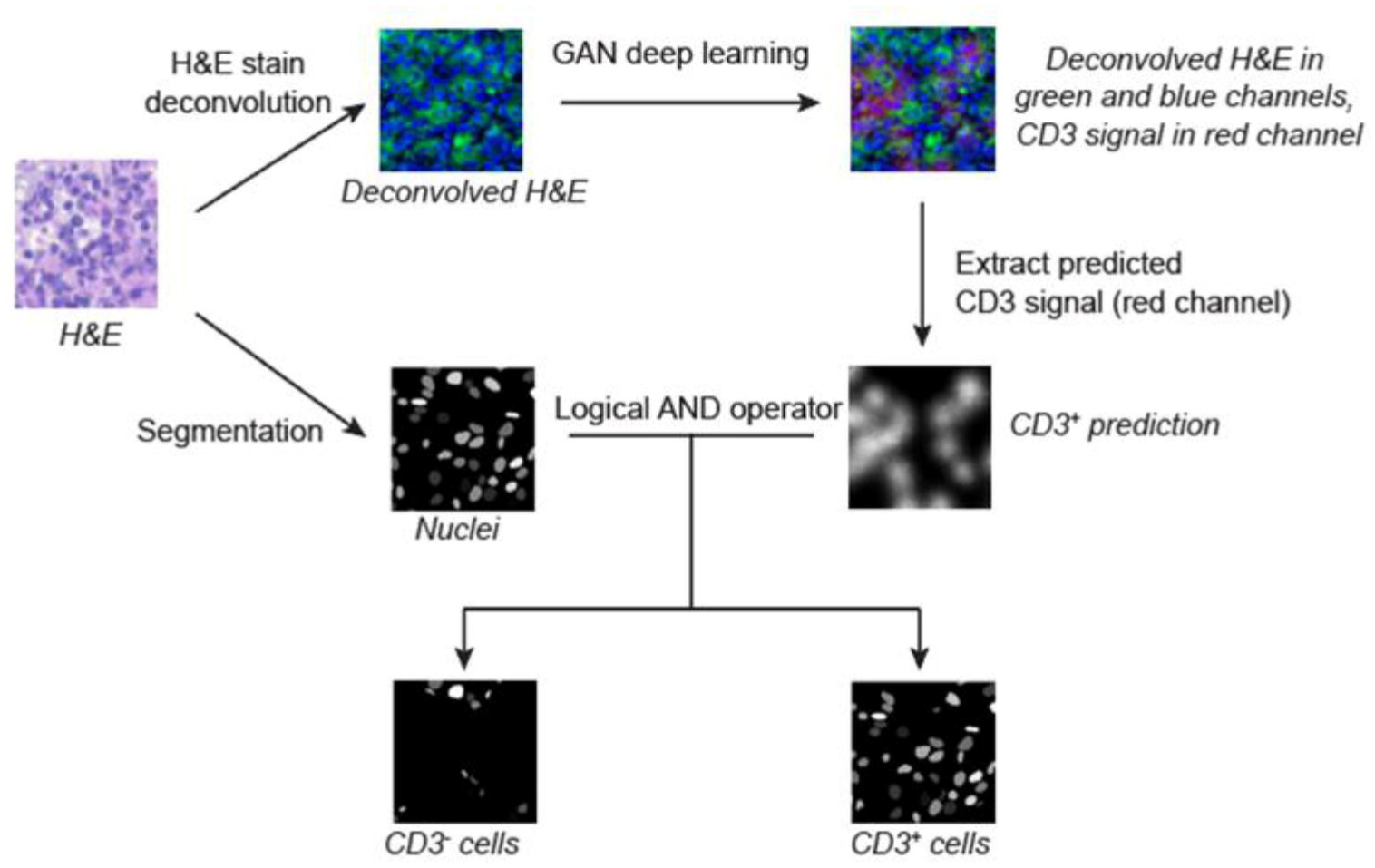
Identification of CD3^+^ and CD3^-^ T-cells predicted by our proposed P2P-GAN models. The process involves extracting the model-predicted CD3 signals (in the red channel) and overlaying the detected signal onto nuclei detected in the H&E image. Nuclei with matching CD3 signals are regarded as CD3^+^ T-cells, otherwise the nuclei are regarded as CD3^-^ T-cells.

## Results

### Validating model performance using training samples

As a sanity check, we assessed the model performance with the image patches used for training (Table 1; N = 57). Of note, the same-section and serial-section datasets were used for testing the same-section and serial-section models, respectively. The predicted CD3^+^ and CD3^-^ T-cell counts from both same-section and serial-section models were highly comparable to the mIF-quantified CD3^+^ and CD3^-^ T-cell counts (i.e., ground truth; all p < 0.005) with Pearson’s correlation >0.95 (Figure 4a-d). However, based on the Mann-Whitney U-test, the same-section model outperformed the serial-section model by a slight margin in terms of overall accuracy (Figure 4e; p < 0.005).

**Figure 4:**
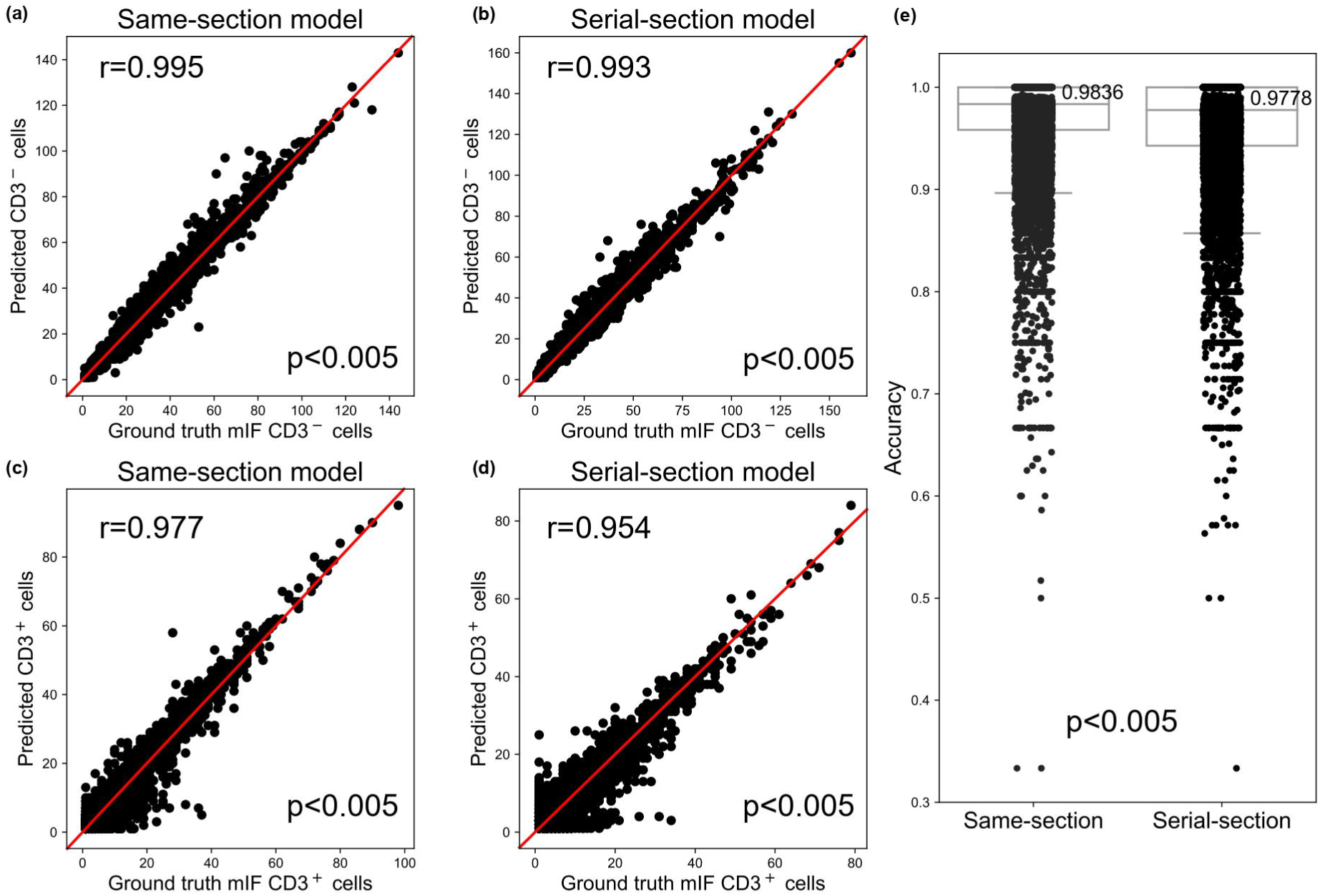
Model performance evaluation using the corresponding training cohorts (i.e., same­ section and serial-section datasets, respectively). Comparison of model-predicted (a-b) CD3^-^ and (c-d) CD3^+^ cell (y-axis) counts with mIF-quantified CD3^+^ cell counts (x-axis) using Pearson’s correlation analysis. (e) Overall accuracy comparison between the model prediction accuracy (y-axis) of the same-section (left) and serial-section (right) models, using the randomly selected held-out samples from the same-section training cohort based on Mann­ Whitney U-tests; each dot represents an image patch.

### Performance comparison of same-section and serial-section models with held-out training cohort

We randomly selected 4% of image patches (384 patches) from the same-section training cohort for model testing. While model-predicted CD3^+^ and CD3^-^ T-cell counts from both the same-section and serial-section models were reasonably comparable to the mIF-quantified CD3^+^ and CD3^-^ T-cell counts (i.e., ground truth) (all p < 0.005, Figure 5), same-section model predictions showed better concordance with the ground truth as compared with that of serial-section model (Pearson’s correlation coefficients 0.784 vs. 0.733, and 0.675 vs. 0.57, respectively; Figure 5a-d). Based on Mann-Whitney U-tests, there was no significant difference in the overall accuracy of the same-section and serial-section models (Figure 5e; p = 0.62).

**Figure 5:**
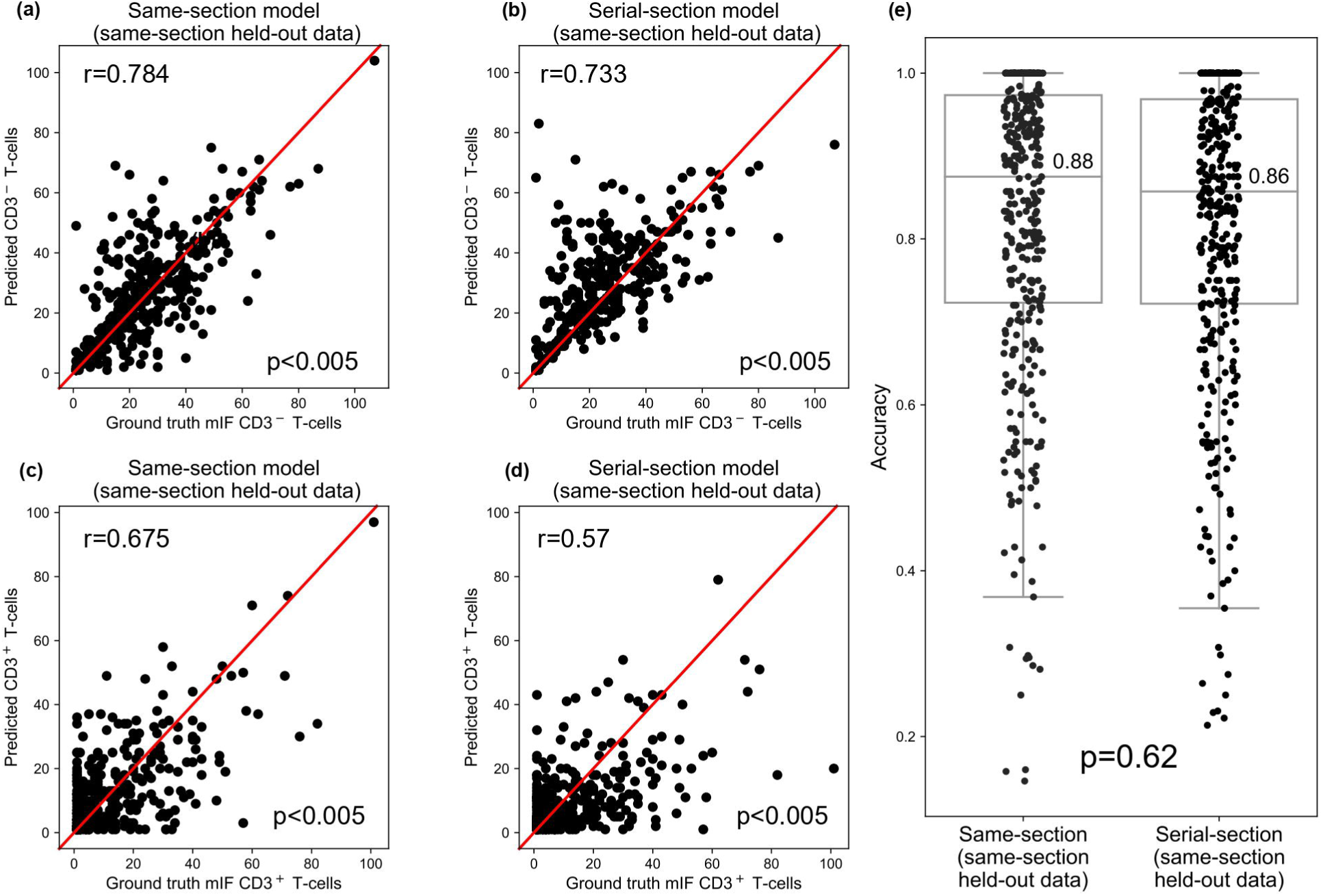
Model performance evaluation using the randomly selected held-out samples from the same-section training cohort. Comparison of model-predicted (a-b) CD3^-^ and (c-d) CD3^+^ cell (y-axis) counts with mIF-quantified CD3^+^ cell counts (x-axis) using Pearson’s correlation analysis. (e) Overall accuracy comparison between the model prediction accuracy (y-axis) of the same-section (left) and serial-section (right) models, using the randomly selected held-out samples from the same-section training cohort based on Mann-Whitney U-tests; each dot represents an image patch.

### Performance comparison of same-section and serial-section models on an independent IHC cohort (N = 48)

In agreement with the results from the held-out cohort analysis, the CD3^+^ and CD3^-^ T-cell counts predicted by both the same-section and serial-section models (Figure 6a) corresponded closely to the IHC-quantified counts, representing the ground truth (p < 0.005, Figures 6b-e). Importantly, the same-section model outperformed the serial-section model, displaying stronger correlations with the IHC ground truth (Figure 6b-c; CD3^-^ T-cell *r* = 0.85 vs. 0.678; Figure 6d-e; CD3^+^ T-cell *r* = 0.886 vs. 0.798), and achieving a higher average accuracy (Figure 6f; mean accuracy = 0.92 vs. 0.65).

**Figure 6:**
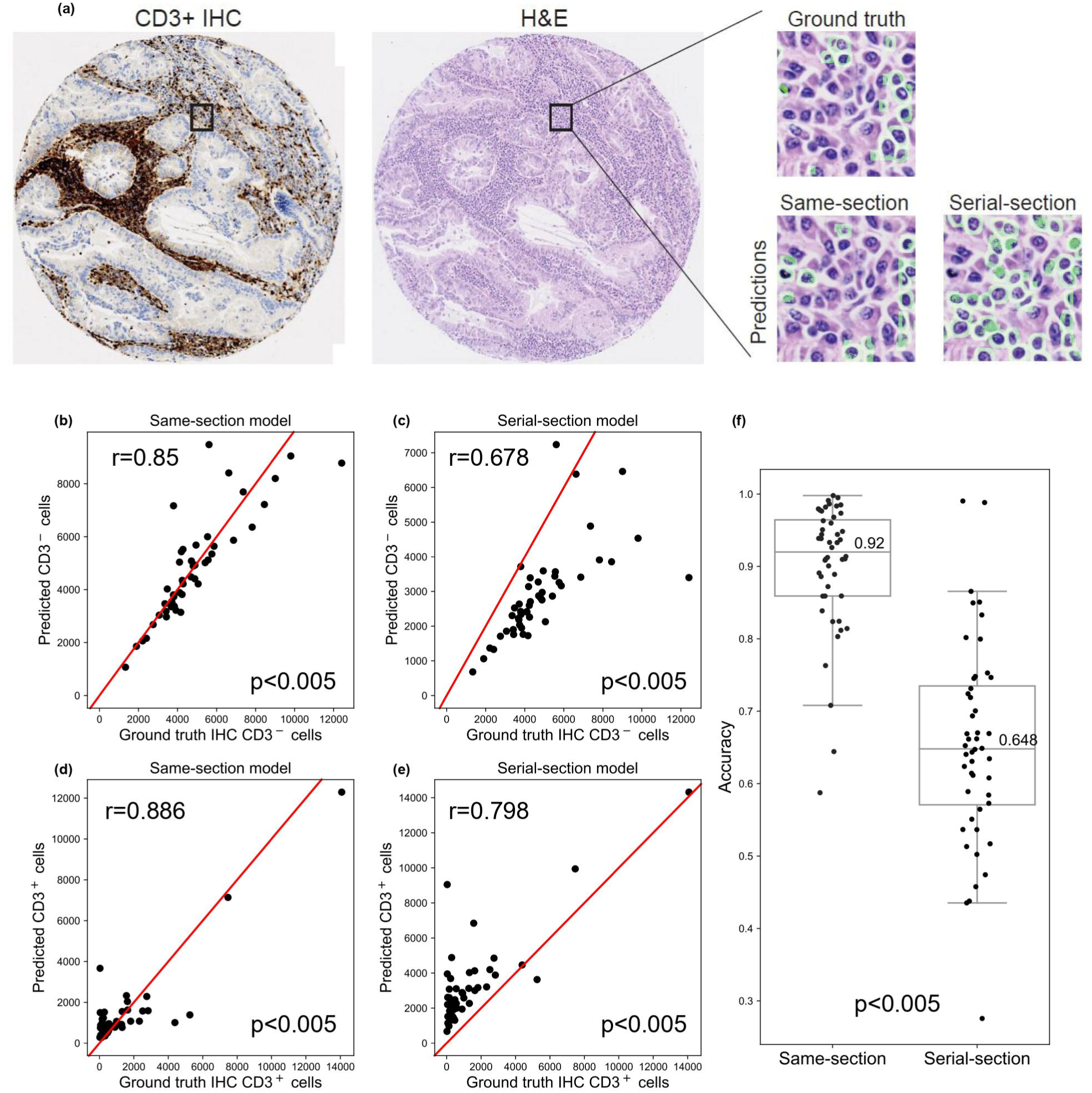
Model performance evaluation using the randomly selected held-out samples from the same-section training cohort with representative images along with model predicted CD3^+^ T-cell visualization in (a). Comparison of model-predicted (b-c) CD3^-^ and (d-e) CD3^+^ (y-axis) cell counts with mIF-quantified CD3^+^ cell counts (x-axis) using Pearson’s correlation. (f) Overall accuracy comparison between the model prediction accuracy (y-axis) of the same-section (left) and serial-section (right) models, using the randomly selected held-out samples from the same-section training cohort based on Mann-Whitney U-tests; each dot represents an image patch.

### Validating the prognostic association of model-predicted CD3^+^ T-cells

Evaluation of the models’ performance on the public Onco-SG cohort (Figure 7a), composed of 204 lung samples (Table 1), revealed a significant correlation between model-predicted CD3 patient groups and 5-year overall survival (Figure 7b; p = 0.013). This association was more pronounced than that observed when patient stratification was based on manual TIL scoring by two pathologists (Figure 7c-d; p = 0.3 and p = 0.06), suggesting the added value of our model in predicting patient outcomes. Nonetheless, the abundance of model-predicted CD3^+^ T-cells showed significant correspondence with the TIL scoring by both pathologists (Figure 7e; p < 0.05).

**Figure 7:**
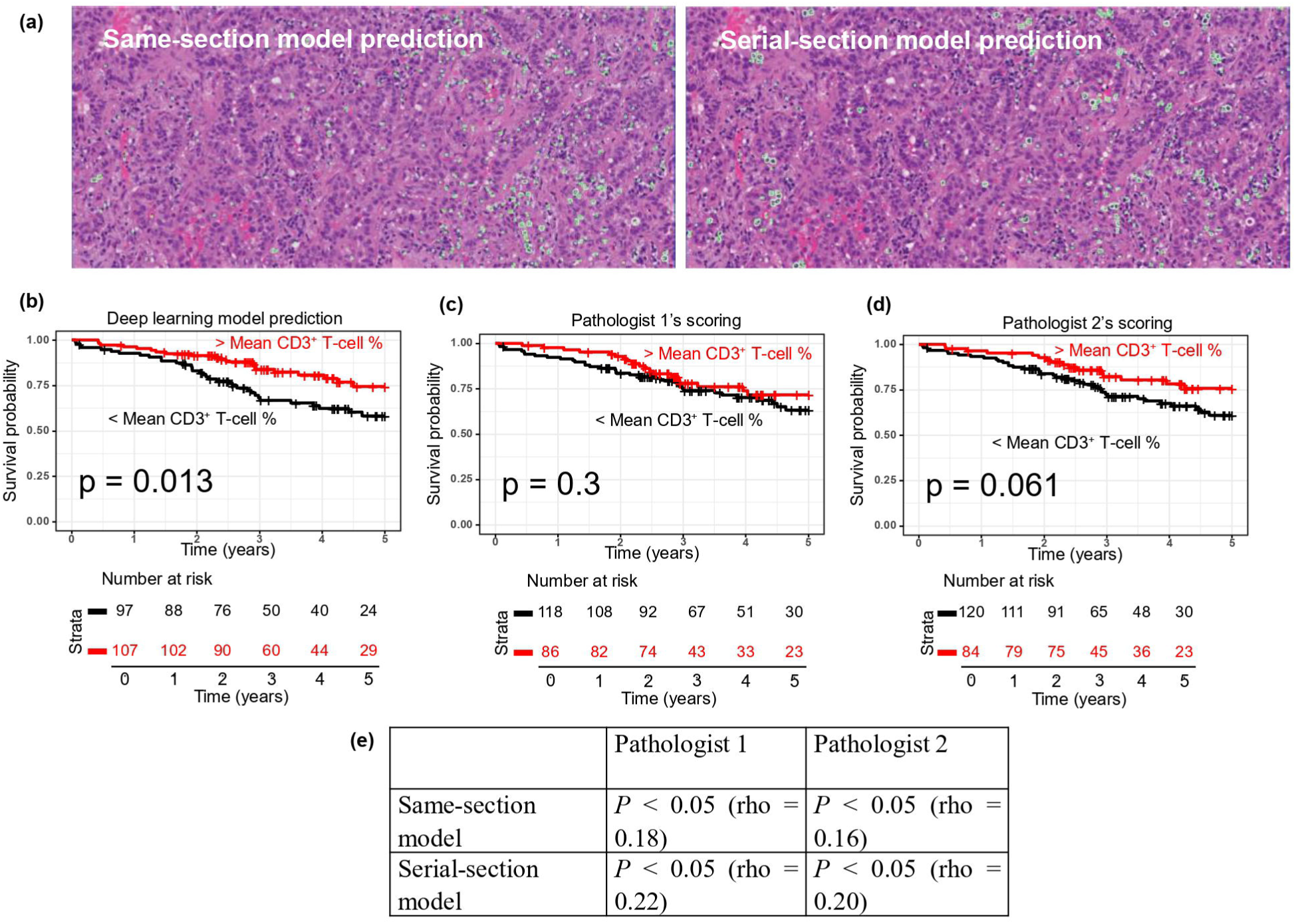
Model performance evaluation using an external lung cohort, with representative images along with model predicted CD3^+^ T-cell visualization in (a). Survival analyses using the external lung cohort show (b) significant association between (same-section) model­predicted CD3 patient groups (low versus high %CD3^+^ abundance groups based on the average %CD3^+^ T-cell counts), while no significant association was observed with the use of manual TIL scoring by (c) pathologist 1, and (d) pathologist 2. (e) Spearman correlation of the prediction of CD3^+^ densities using the same- and serial-section models with the manual TIL density scoring by two independent pathologists.

## Discussion

In this study, we developed and examined P2P-GAN virtual staining models to predict CD3^+^ T-cells from low-cost digitized H&E images. A significant aspect of our investigation was the exploration of performance disparities that arise when ground truth cell labels are obtained from the same tissue section used for H&E staining, as opposed to a serial section. Our findings demonstrate that the model trained using the same-section approach consistently surpasses the serial-section model. This superiority manifests as stronger correlations with mIF and IHC-quantified CD3^+^ and CD3^-^ T-cells, along with heightened overall prediction accuracies. It also reinforces the potential of the same-section model as a robust technique in histopathology-driven immune phenotyping. Crucially, our work also showcased the enhanced prognostic utility of our model-predicted CD3^+^ T-cell abundance when compared to traditional manual TIL scores. This emphasizes the clinical relevance of our proposed virtual staining model in a real-world setting, potentially facilitating improved patient stratification and treatment decision-making. A distinctive feature that sets our proposed model apart from traditional DL models for cell prediction is its capability for image-to-image translation, virtually staining the CD3 marker within the original H&E context. This has two major implications. First, it facilitates further downstream analysis of the TME and spatial interplay between predicted cell types and other cellular or tissue data derived from H&E through either pathological assessment or digital pathology. Second, it creates a new pathway for integrating incremental cell type predictions from different models onto the same H&E space. Collectively, these advancements could significantly enhance our understanding of the TME, potentially leading to the identification of novel spatial biomarkers or therapeutic targets.

While our proposed approach has yielded encouraging results, it is important to acknowledge its inherent limitations. First, our current model is designed specifically for CD3^+^ T-cells prediction from H&E images and may not generalize well to other cell types or markers without significant adjustments or retraining. Additionally, its performance may be compromised when applied to tumor types beyond lung cancer. Second, the application of this model is largely limited to high-quality digital slides. Its performance may be affected by variations in tissue preparation, staining procedures, and image acquisition methods across different laboratories. Nevertheless, the clinical significance of our model has been validated using a publicly available lung cohort. Lastly, despite overall robust performance, we noted outliers in our model’s predictions, indicating potential areas for improvement. These discrepancies suggest complex, unaddressed variables within biological samples that need further investigation. Future endeavors should focus on understanding these outlier causes, refining modeling techniques, and incorporating larger, more diverse datasets for improved generalizability and outlier management.

In conclusion, our thorough exploration into the necessity of employing ground truth cell labels from identical tissue sections in a CD3^+^ T-cell prediction model signifies a notable advance in the domain of H&E-based virtual staining research. Our novel image-to-image translation capability paves the way for in-depth TME analyses. Combined with the potential of predicting refined cell types via the mIF technique, our model unveils exciting new possibilities for biomarker discovery and the advancement of therapeutic strategies. While certain limitations are observed, these challenges underscore the direction for future investigations, the results of which could greatly enhance the prediction accuracy and clinical applicability of this innovative approach.

## Acknowledgements

This work was supported by the Bioinformatics Institute (BII), the Singapore Immunology Network (SIgN), the Institute of Molecular and Cell Biology (IMCB) and the Agency for Science, Technology and Research (A*STAR).

## Ethics Approval and Consent to Participate

This study was approved by the Agency of Science, Technology and Research (A*STAR) Human Biomedical Research Office (A*STAR IRB: 2021-161, 2021-188, 2021-112).

## Author Contributions

J.P.S.Y, M.C.L. and Y.C. conceived and directed the study. A.B.A. performed the development, training and testing of the DL models and conducted the biostatistical analysis; B.L.C. performed the testing of codes. F.W. and J.C.T.L. performed immunohistochemical techniques; J.P.V. and

Y.Z.X. performed the TIL scoring. W.W.Y. created the publication figures. D.S.W.T, A.T., C.C.Y., L.Y.K. and T.K.H.L. conducted the sample acquisition and provided clinical pathological and oncological perspectives. A.B.A., F.W. and M.C.L. prepared the manuscript. All authors reviewed the manuscript.

## Funding

We would like to express our gratitude towards the following parties for their valuable contributions to this work:

A*STAR BIOMEDICAL ENGINEERING PROGRAMME (Project No: C211318003), Singapore National Medical Research Council (MOH-000323-00, OFYIRG19may-0007), IAF-PP (HBMS Domain): H19/01/a0/024-SInGapore ImmuNogrAm for Immuno-Oncology (SIGNAL)

Bioinformatics Institute and Singapore Immunology Network, Agency for Science, Technology and Research (A*STAR), A*STAR Biomedical Engineering Programme (C211318003)

Industry Alignment Fund-Industry Collaboration Fund (IAF-ICP I2201E0014)

Singapore National Medical Research Council (MOH-000323-00, OFYIRG19may-0007).

## Data Availability Statement

The mIF and in-house IHC data sets used during the current study are available from the corresponding author upon reasonable request. The external lung cancer cohort is available in the OncoSG repository, https://src.gisapps.org/OncoSG/. The scripts used in this study can be found in the following GitHub repository, https://github.com/abubakrazam/Pix2Pix_TIL_H-E.git

